# Sensory Neuropeptides Dictate Sex-Specific Synovial Immunity and Cartilage Degeneration in Aging Mice

**DOI:** 10.64898/2026.07.27.740931

**Authors:** Patrick Pann, Ravikumar Mayakrishnan, Babak Moradi, Brian Johnstone, Susanne Grässel

**Affiliations:** Dept. of Orthopedic Surgery, Experimental Orthopedics, Center for Medical Biotechnology, Bio Park 1, University of Regensburg, Germany; Orthopedic Research Center, Kiel University, Germany; Department of Orthopedics and Trauma Surgery, University Medical Center Schleswig-Holstein, Campus Kiel, Germany; Department of Orthopedics and Rehabilitation, Oregon Health & Science University, Portland, Oregon, USA

**Keywords:** aging, substance P, alpha-calcitonin gene-related peptide, articular cartilage, synovia

## Abstract

Sensory neuropeptides, particularly Substance P (SP) and α-calcitonin gene-related peptide (αCGRP), are implicated in osteoarthritis (OA) pathogenesis. This study elucidates their specific roles in spontaneous, age-related OA. Male and female mice deficient in SP (*Tac1*−/−), αCGRP (αCGRP−/−), or both (DKO) were evaluated at 6, 12, and 18 months of age. Assessments included histological OARSI scoring for articular cartilage matrix structure, Luminex arrays for systemic serum cytokines, and flow cytometry for local synovial immune cell profiling. Wild type (WT) mice developed early-stage, age-related cartilage degradation, predominantly in the lateral compartment. Conversely, all neuropeptide-deficient strains exhibited significant structural protection against this process. Systemically, SP deficiency distinctly altered cytokine profiles (e.g., decreased IL-23, increased IP-10), whereas αCGRP deficiency caused minimal systemic shifts, highlighting a disconnect between circulating markers and local joint preservation. Locally, flow cytometry revealed profound, sexually dimorphic, and age-dependent neuroimmune alterations. In young males, neuropeptide deficiency significantly reduced synovial macrophage counts to levels comparable to those of aged WT mice. Furthermore, male αCGRP−/− mice exhibited an age-related accumulation of CD8+ cytotoxic T cells. In contrast to males, young WT females demonstrated higher baseline CD8+ T cell counts that declined with age, whereas these subpopulations remained persistently low in KO mice. SP and αCGRP act as critical modulators of age-related cartilage degradation. Their absence provides robust structural protection mediated through highly localized, sexually dimorphic neuroimmune pathways. These findings emphasize the necessity of targeting the local joint microenvironment for future personalized, sex-specific OA therapies.

## Introduction

Aging is one of the strongest risk factors for osteoarthritis (OA) development which is a degenerative whole-joint disease mainly characterized by articular cartilage breakdown, subchondral bone remodeling, and synovial inflammation. The peripheral sensory nervous system plays a fundamental role in regulating this joint tissue homeostasis during aging and OA pathogenesis besides pain transmission [1–3]. The sensory neuropeptides substance P (SP) and alpha-calcitonin gene-related peptide (αCGRP) function as key neuro-signaling molecules modulating these processes. While SP and αCGRP are released from afferent nerve terminals innervating joint tissues, they are also endogenously synthesized and secreted by resident joint cells, including synoviocytes, osteoblasts, osteoclasts, and chondrocytes [1, 2]. This local production allows SP and αCGRP to regulate cartilage, synovial and bone metabolism directly through autocrine and paracrine mechanisms [2].

Physiological aging is a primary driver of OA, and both SP and αCGRP significantly influence age-related joint deterioration [4]. Murine genetic knockout models have demonstrated the individual contributions of these neuropeptides to OA pathology. Mice lacking αCGRP (encoded by the *Calca* gene) exhibit protection against age-dependent cartilage degradation and joint inflammation but develop severe subchondral bone sclerosis, indicating a dual pro-inflammatory and bone-protective role [4, 5]. Similarly, deficiency in SP (encoded by the Tachykinin precursor 1 gene; *TAC1*) alters cartilage matrix stiffness, impacts cartilage preservation, and promotes subchondral bone alterations following mechanical joint destabilization [6, 7]. Despite their known individual roles and frequent co-expression in sensory nerves, the synergistic effects of SP and αCGRP on age-related joint tissue degradation ultimately leading to OA remain not well defined [3, 8].

We hypothesize that SP and αCGRP play an interactive role in driving the pathogenesis of age-related OA. To test this hypothesis, the present study longitudinally analyzes male and female wild type (WT) mice alongside single and double genetic knockouts for SP and αCGRP. These animal models are evaluated across distinct chronological stages at 6, 12, and 18 months of age. The study quantifies cartilage degradation using the OARSI histopathological grading score, characterizes shifts in synovial tissue immune cell populations, and profiles OA-associated and proinflammatory serum markers. This comprehensive approach aims to define the specific neuroimmune mechanisms by which these neuropeptides interactively influence articular tissue homeostasis during aging.

## Methods

### Animals

The housing conditions and origins of the mice were identical to those detailed in our previous study [9]. The mice were maintained under a 12-hour light/dark cycle with ad libitum access to both food and water (Ethical Approval number: AZ 55.2-2532-2-1253). For control purposes, 8-to 10-week-old wild type (WT) C57Bl/6J mice were obtained from Charles River Laboratories (Sulzfeld, Germany). The tachykinin 1 knockout strain (Tac1−/−, substance P-deficient) was kindly provided by A. Zimmer from the University of Bonn [10]. The alpha-calcitonin gene-related peptide-deficient line (αCGRP−/−), engineered by R.B. Emeson [11] was a generous gift from T. Schinke and M. Amling (University Medical Center Hamburg-Eppendorf). Both mutant strains had been backcrossed onto a C57Bl/6J genetic background. A double knockout (DKO) strain was generated by cross-breeding Tac1−/− and αCGRP−/− mice. The experimental cohort comprised a total of 67 WT, 42 Tac1−/−, 45 αCGRP−/−, and 41 DKO animals. Mice were euthanized via carbon dioxide inhalation at the ages of 6, 12, and 18 months and blood and knee joints were removed.

### Serum Analysis

Immediately following euthanasia, blood samples from 8, 12 and 18 months old mice were collected via intracardiac puncture. Serum was separated by centrifugation to quantify pro-inflammatory and OA-associated factors using multiplex immunoassays. The analysis was performed on a Bio-Plex 200 Multiplex Reader (Bio-Rad Laboratories GmbH) using a Mouse ProcartaPlex Mix&Match 13-plex assay (PPX-13-MXAAD4K, Thermo Fisher Scientific). This panel comprised the following cytokines and chemokines: GM-CSF, IL-10, IL-15/IL-15R, IL-17A (CTLA-8), IL-23, IL-4, IP-10 (CXCL10), MCP-1 (CCL2), MIG (CXCL9), MIP-1α (CCL3), MIP-1β (CCL4), RANTES (CCL5), and VEGF-A. Analytes that failed to reach the assay limit of quantification were excluded from further analysis and are omitted from the results and supplementary data.

### Histology and OA Scoring

Histological preparation and OA grading followed the methodologies previously established in [9]. Upon dissection, knee joints from 8, 12 and 18 months old mice were fixed in 4% paraformaldehyde/PBS for 24 hours and subsequently decalcified in 20% EDTA (pH 7.4; Carl Roth) over a period of eight weeks. Following paraffin embedding, a microtome was used to cut 6-μm frontal sections. To evaluate cartilage degradation, a total of 12 sections—comprising six pairs of consecutive sections spaced 60–90 μm apart—were deparaffinized, rehydrated, and stained with Safranin O (Sigma-Aldrich), Weigert’s iron hematoxylin (Carl Roth, Merk), and Fast Green (PanReac Applichem). Two independent, blinded investigators evaluated the specimen sections according to the OARSI guidelines [12]. Digital micrographs were captured at 10× magnification utilizing a BZ-X810 microscope (KEYENCE Deutschland GmbH). Quantification was based on the calculated mean OARSI scores from both the lateral and medial femoral condyles and tibial plateaus.

### FACS Analysis

For tissue processing, synovial samples pooled from 4 to 5 mice at 6 and 18 months of age were finely minced and enzymatically digested with Liberase TL (Sigma-Aldrich, 1 U/mL) for 60 min at 37 °C. The resulting cell suspension was passed through a 70–75 μm mesh, pelleted by centrifugation at 300×g for 10 min, and resuspended in autoMACS® Running Buffer [13].

For immunophenotyping, cell-surface antigens were labeled by adding 2 μL of each fluorochrome-conjugated antibody to 100 μL of the cell suspension in staining buffer. For the lymphoid surface panel, antibodies against CD3 (#130-119-793), CD4 (#130-123-894), CD8a (#130-123-240), and CD25 (#130-126-615) (all Miltenyi Biotec) were added; for the myeloid panel, the surface cocktail comprised antibodies against CD11b (#130-113-810), Ly6G (#130-123-029), F4/80 (#130-118-459), Ly6C (#130-111-919), CD11c (#130-110-838) (all Miltenyi Biotec), and CD206 (BioLegend, #Biolegend 141721). Viable cells were distinguished from dead cells using Viobility™ Fixable Dye (Miltenyi Biotec, Bergisch Gladbach, Germany; Cat# 130-130-421) following the manufacturer’s protocol. After a 10-min incubation at 4 °C in the dark, samples were diluted with 1 mL of autoMACS buffer, centrifuged at 300×g for 10 min, and resuspended in 100 μL of fresh autoMACS buffer. To analyze intracellular targets within the lymphoid panel, the surface-labeled cells were subsequently fixed using the Fixation Buffer from the FoxP3 Staining Buffer Set (Miltenyi Biotec; # 130-093-142) for 30 min at 4 °C. Following a wash step, cells were permeabilized with Permeabilization Buffer for 10 min at 4 °C. Intracellular staining was performed by incubating the cells with antibodies against IFN-γ (#130-117-502), IL-17A (#130-112-008), FoxP3 (#130-111-682) (all Miltenyi Biotec), and IL-4 (BioLegend, # Biolegend 504125) (2 μL each) for 30 min at 4 °C in the dark. Samples were then washed with autoMACS buffer, centrifuged, and resuspended for data collection.

Data acquisition was performed on a MACSQuant® 16 flow cytometer (Miltenyi Biotec) utilizing standard compensation matrices. The sequential gating strategy progressed from total cells to singlets, and subsequently to live cells, before lineage-specific analyses were executed (**Suppl. Fig. 1**). Within the lymphoid compartment, total T cells were identified as CD3+ cells, which were further subdivided into helper T cells (CD3+CD4+) and cytotoxic T cells (CD3+CD8+). The CD3+CD4+ population was subsequently gated to evaluate T helper cell subsets using CD25, FoxP3, IFN-γ, IL-17A, and IL-4 expression. For myeloid lineage analysis, gates were restricted to non-lymphoid live singlets, defining total myeloid cells as the CD11b+ population. Within this compartment, neutrophils were characterized as CD11b+Ly6G+, and tissue macrophages were identified as CD11b+F4/80+Ly6C^low. Differential expression of Ly6C was utilized to separate these macrophages from Ly6C^high monocytes. Finally, tissue macrophages were evaluated for CD11c and CD206 expression to characterize M1-and M2-activated subpopulations. All flow cytometry data were processed using FlowJo software (BD Bioscience), with cell frequencies expressed as percentages of their respective live-cell parent gates [13].

### Statistical Analysis

Statistical processing and data visualization were executed using GraphPad Prism version 11.0.2 (San Diego, CA, USA). Outliers within the OARSI scores and serum datasets were identified via the ROUT method, utilizing thresholds of Q = 0.1 % and Q = 1 %, respectively [14]. For FACS data, outlier detection was performed using Grubbs’ test with a significance level of alpha = 0.01. To evaluate the effects of genotype and age across groups, a two-way ANOVA with Tukey’s post hoc tests for multiple comparisons or a Kruskal-Wallis with Dunn’s test for multiple comparisons was conducted. Graphical data are depicted as box plots, where the horizontal lines represent the medians, the boxes indicate the interquartile ranges, and the whiskers extend to the minimum and maximum values.

## Results

### Substance P deficiency drives systemic elevation of IP-10, MIG, and RANTES alongside IL-23 suppression

To evaluate systemic inflammatory profiles, serum concentrations of aging/OA-associated cytokines and chemokines were quantified across all experimental groups (**Fig. 1**). Male and female Tac1−/− and DKO (Tac1−/− αCGRP−/−) mice exhibited significantly elevated serum concentrations of IP-10 (CXCL10), MIG (CXCL9), and RANTES (CCL5) compared to WT mice across all investigated age groups. In contrast, the αCGRP−/− strain demonstrated no significant deviations from WT baseline levels for any of these markers. While no general age-dependent alterations were observed for these chemokines, a pronounced sex-specific variance was detected exclusively in RANTES. Female cohorts presented with approximately five-fold higher systemic RANTES concentrations compared to their male counterparts across the respective genotypes.

**Figure 1.**
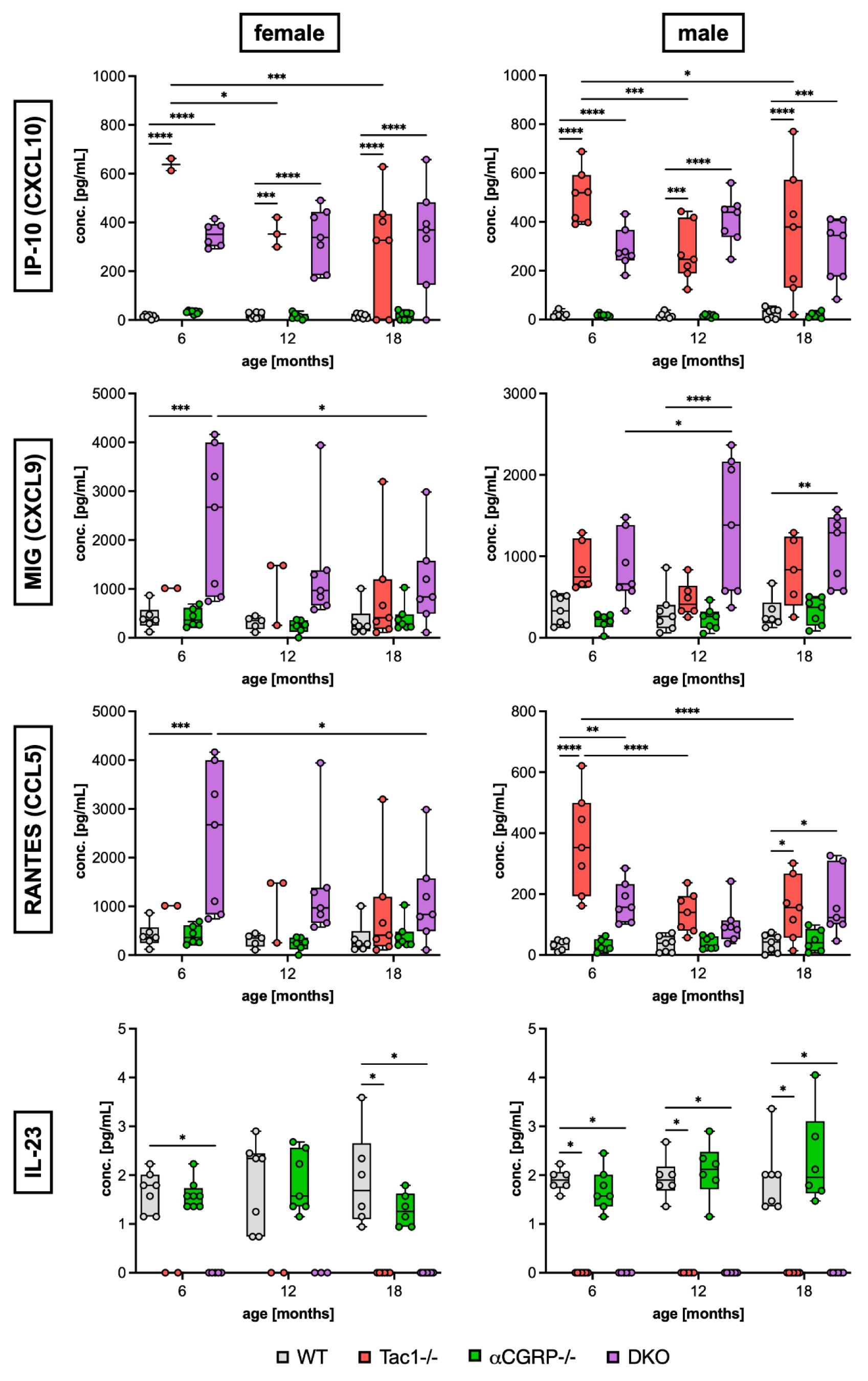
Systemic profiling of osteoarthritis-associated chemokines and cytokines across aging neuropeptide-deficient mice. Serum marker concentrations of female (left) and male (right) WT and KO mice strains at 6, 12, and 18 months of age. Statistical analysis is using two-way ANOVA with Tukey’s test for multiple comparisons (IP-10, MIG, RANTES) or Kruskal-Wallis with Dunn’s test for multiple comparisons (IL-23). * p < 0.05, ** p < 0.01, *** p < 0.001, **** p < 0.0001. N = 2-8.

In direct opposition to the observed chemokine elevations, serum concentrations of IL-23 were profoundly diminished in both male and female Tac1−/− and DKO mice. Across all evaluated ages, IL-23 concentrations in these specific strains dropped below the lower limit of quantification. Consistent with the data described above, αCGRP−/− mice maintained IL-23 levels comparable to WT animals, and no systemic age-or sex-dependent fluctuations were detectable.

Due to a high number of individual samples dropping below the quantification limit, reliable data is not available for the serum markers MIP-1β (CCL4), GM-CSF, and VEGF-A (**Suppl. Fig. 2**). MIP-1ß concentrations were mostly similar across all groups and GM-CSF concentrations were found to be reduced across neuropeptide-deficient strains compared to WT mice, alongside sporadic, age-independent VEGF-A elevations primarily within the Tac1−/− cohorts. However, due to the compromised data quality, these supplementary parameters lack the robustness and consistency required for definitive conclusions.

### Sensory neuropeptide deficiency induces sex-and age-specific shifts in synovial leukocyte populations

To characterize the local inflammatory milieu, synovial leukocyte populations from mouse knee joints were quantified using flow cytometry (**Fig. 2**). The comprehensive analysis revealed profound sex-, age-, and genotype-dependent alterations across both lymphoid and myeloid lineages.

**Figure 2.**
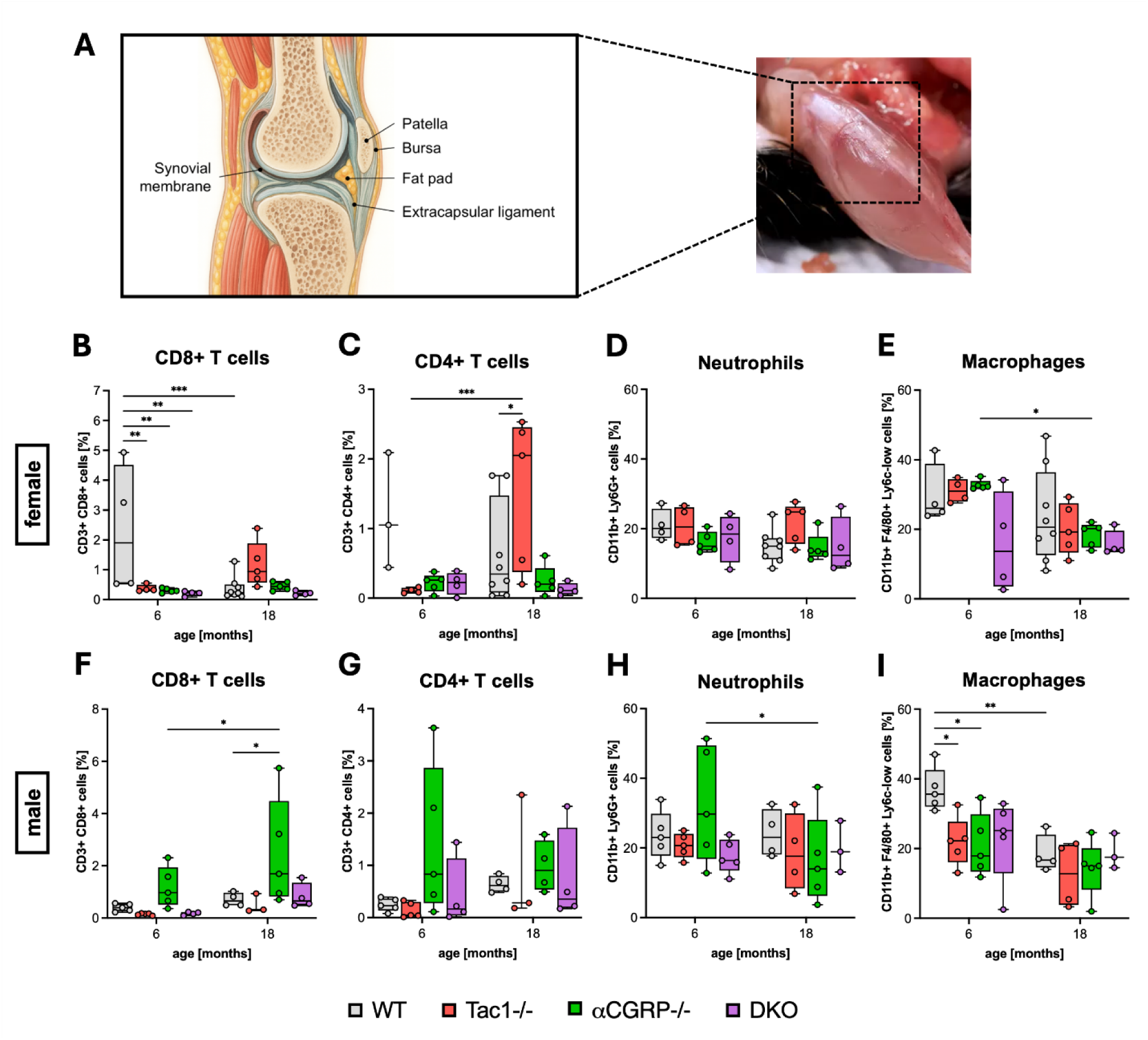
Flow cytometric characterization of major synovial myeloid and lymphoid leukocyte populations during joint aging. Mouse knee synovial tissues were dissected and analyzed by flow cytometry. Flow cytometry analysis shows the percentages of different synovial leukocyte populations in female (B-E) and male (F-I) WT and KO mice at 6 and 18 months of age. Statistical analysis is using two-way ANOVA with Tukey’s test for multiple comparisons. * p < 0.05, ** p < 0.01, *** p < 0.001. N = 3-8.

Evaluation of the cytotoxic T cell compartment (CD8+) demonstrated that 6-month-old female WT mice possessed significantly higher cell frequencies compared to age-matched KO strains. By 18 months of age, these CD8+ levels declined, aligning with the lower frequencies observed in the respective KO groups (**Fig. 2B**). Conversely, male αCGRP−/− mice exhibited a significant age-dependent expansion of CD8+ cells at 18 months, representing a significant increase both relative to their 6-month baseline and compared to 18-month-old WT controls (**Fig. 2F**). Within the helper T cell (CD4+) population, female Tac1−/− mice demonstrated a significant accumulation of CD4+ cells at 18 months of age, reaching levels significantly higher than both their 6-month baseline and the 18-month-old WT animals (**Fig. 2C**). In contrast, male mice displayed no significant age-or genotype-dependent variations (**Fig. 2G**). Synovial neutrophil frequencies remained largely consistent across both sexes and most genotypes (**Fig. 2D, H**). A notable exception occurred in male αCGRP−/− mice, which experienced a significant concurrent reduction in neutrophils at 18 months of age compared to the 6-month time point. Macrophage populations exhibited distinct, sex-specific temporal dynamics (**Fig. 2E, I**). In female cohorts, αCGRP−/− mice showed a significant age-related decline in macrophages from 6 to 18 months, a trend that was similarly observed in female WT and Tac1−/− mice. Female DKO mice presented with lower baseline macrophage levels already at 6 months, which remained unchanged at 18 months. In male cohorts, all three KO strains exhibited significantly reduced macrophage frequencies at 6 months of age relative to WT mice. By 18 months, macrophage levels in male WT mice decreased, eliminating the genotypic differences at a younger age.

Due to high standard deviations, data for specific T cell-and macrophage subpopulations lack the robustness and consistency required for definitive conclusions (**Suppl. Fig. 3**). Under this premise, the data allow the hypothesis of an αCGRP-driven effect on Th1 polarization, given that female αCGRP−/− and DKO mice exhibited a significant increase in Th1 cells at 18 months relative to 6 months. In contrast, Th1 frequencies in female WT and Tac1−/− mice appeared to decline over the same period (**Suppl. Fig. 3A, B**). Furthermore, the data suggest the possibility of a SP-dependent influence, indicated by a significant reduction in Th2 frequencies in 18-month-old female Tac1−/− mice compared to their 6-month baseline (**Suppl. Fig. 3C**). In all other T cell-and macrophage subpopulations no significant differences between mouse strains, ages or sexes were detectable (**Suppl. Fig. 3D-L**).

### Sensory neuropeptide deficiency protects against age-associated lateral cartilage degradation

To assess the influence of age, sex, and the absence of sensory neuropeptides on articular cartilage integrity, the knee joints of WT, single KO and DKO mice at 6, 12, and 18 months of age were evaluated histologically using the OARSI scoring system (**Fig. 3**).

**Figure 3.**
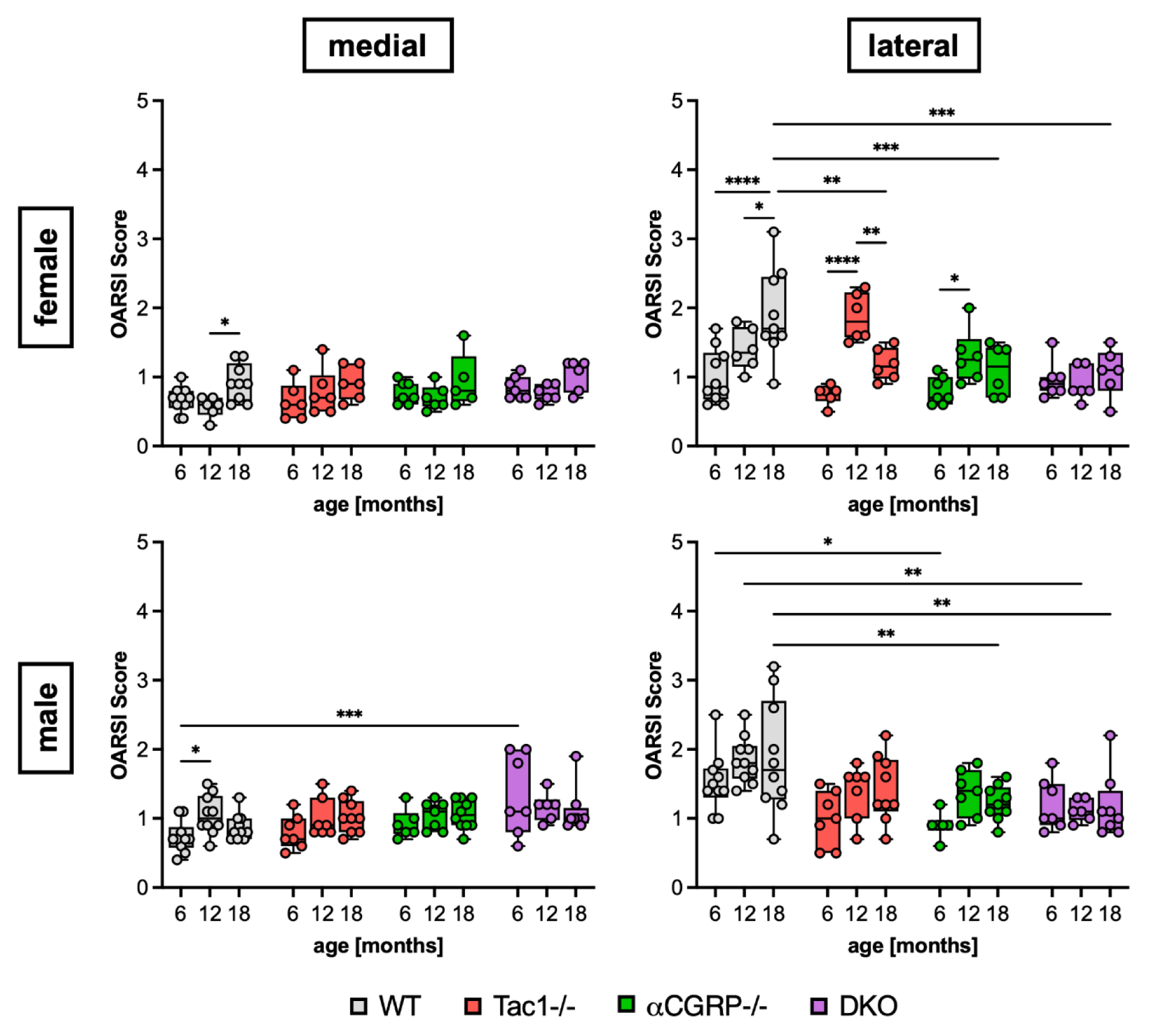
Histological evaluation of age-and genotype-dependent morphological joint destruction in neuropeptide-deficient mice. Cartilage of the right knee joints of female and male WT and KO mice at ages 6 and18 months was evaluated for grades of destruction according to the OARSI guidelines for murine OA. Mean OARSI scores of the medial and lateral sides were calculated from both tibial and femoral scores. Statistical analysis is using two-way ANOVA with Tukey’s test for multiple comparisons. * p < 0.05, ** p < 0.01, *** p < 0.001. N = 6–10.

Overall, both male and female mice exhibited relatively low levels of cartilage matrix degradation on the medial side across all investigated genotypes, with mean OARSI scores predominantly ranging between 0.5 and 1.5. Significant age-related increases relative to the 6-month baseline were restricted to specific WT cohorts (18-month-old female WT and 12-month-old male WT mice). Although male DKO mice displayed significantly higher medial scores than age-matched WT animals at 6 months, this effect appears to be driven primarily by high intra-group data variability and did not persist at later time points.

In contrast to the medial side, the most severe cartilage matrix degradation was observed in the lateral compartment. Female WT mice demonstrated a significant, progressive increase in lateral OARSI scores from 6 to 18 months of age. By 18 months, the lateral scores of female WT animals were significantly elevated compared to all three KO strains at the same time point. While female Tac1−/− mice exhibited a significant transient increase in lateral cartilage degradation at 12 months, the scores reverted to baseline levels by 18 months.

In male cohorts, WT mice trended toward higher lateral OARSI scores across the entire investigated age range compared to the neuropeptide-deficient strains. This attenuated cartilage matrix degradation in the KO models reached statistical significance at multiple time points: lateral scores in male WT mice were significantly higher than those in αCGRP−/− mice at 6 and 18 months, and significantly higher than those in DKO mice at 12 and 18 months.

## Discussion

The onset and progression of OA are fundamentally influenced by both age and sex [15–17]. Sensory neuropeptides, specifically SP and αCGRP, have been previously identified as crucial modulators of cartilage and subchondral bone matrix structure in both post-traumatic and age-related joint degeneration [4–6, 9]. To further elucidate this mechanism, the present study investigated the complex interplay of these neuropeptides in spontaneous, age-related osteoarthritis-like processes, utilizing single and double knockout murine models evaluated comprehensively between 6 and 18 months of age.

Our histological analysis demonstrated that male and female WT mice exhibit progressive, age-related cartilage degradation indicative of an early OA phenotype, whereas the KO models were significantly protected. By 18 months of age, the neuropeptide-deficient strains presented with substantially lower OARSI scores compared to their WT counterparts. This structural deterioration was predominantly localized to the lateral compartment. Current literature regarding compartmentalization in age-related murine OA remains ambiguous with reports of equivalent degradation across both compartments or higher medial degradation at similar ages [4, 18, 19]. Furthermore, male subjects are frequently noted to follow a more severe disease progression [4, 18, 19].

The structural protection conferred by SP deficiency in our study aligns with established post-traumatic models, where SP deficiency actively delays cartilage degradation and prevents the stiffening of the extracellular cartilage matrix [6, 7, 9]. Conversely, the role of αCGRP appears to be highly dependent on the local joint environment. Previous post-traumatic models evaluating αCGRP deficiency have yielded conflicting results, with cartilage degradation reported as unaffected, temporarily delayed, or even slightly accelerated [5, 7, 9, 20]. In line with our findings female αCGRP-/-mice were reported to be protected from spontaneous, age-related joint degeneration [4]. These discrepant findings underscore diverging roles for sensory neuropeptides depending on the pathophysiological context, specifically differentiating between acute trauma, active inflammation, and physiological senescence.

On a systemic level, SP deficiency triggered significant alterations in chemokine profiles. Both male and female Tac1−/− and DKO mice exhibited markedly increased serum concentrations of IP-10 (CXCL10), MIG (CXCL9), and RANTES (CCL5), alongside diminished levels of IL-23, when compared to WT and αCGRP-/-mice. The reduction in IL-23 provides a direct mechanistic explanation for the preserved joint structure observed in these strains. IL-23 is strongly associated with OA-related pain in both human patients and murine models, and the targeted deletion of its IL-23p19 subunit is known to protect against chemically induced cartilage degradation and joint inflammation [21–23]. While IL-23 canonically promotes Th17 cell differentiation, our local synovial data only reflected a trend toward Th17 reduction in 18-month-old male WT-and SP-deficient mice, highlighting a potential divergence between systemic cytokine availability and localized cellular responses [24].

Both MIG and IP-10 signal via the CXCR3 receptor to recruit Th1 helper cells [25]. Despite the systemic elevation of these chemokines, significant local increases in synovial Th1 cells were restricted to αCGRP-/-and DKO mice at 18 months of age. While circulating levels of MIG and IP-10 are reported to naturally increase with age in both healthy individuals and OA patients, serum levels remained stable in our WT mice [26]. MIG and IP-10 concentrations even declined slightly with age in Tac1−/− and DKO mice while remaining significantly elevated in comparison to WT mice over the observed timeframe. The local effects of MIG and IL-10 described in the literature differ significantly from one another. Elevated localized MIG is generally linked to synovitis, IL1b expression, and poorer knee function [25, 27]. However, IL-10 is upregulated following joint fracture in mice and has demonstrated the capacity to inhibit total matrix metalloproteinase and aggrecanase activity *in vitro* [28].

The elevated levels of RANTES further complicate the systemic profile. RANTES typically induces catabolic enzymes, including MMP-1 and MMP-13, within synovial fibroblasts, and inhibiting the expression of its primary receptor, CCR5, effectively reduces cartilage degradation in post-traumatic models [29, 30]. Although RANTES – together with MIG and IP-10 – is classically secreted by pro-inflammatory M1 macrophages, our flow cytometry analysis revealed no significant differences in M1 macrophage populations between Tac1−/− and WT mice [31]. Another source can be resident synovial fibroblasts, which are known to highly express RANTES and subsequently release IL-6 upon stimulation [31].

Ultimately, while the combination of reduced IL-23 and elevated IP-10 strongly supports the protection from age-related OA in Tac1−/− and DKO mice, the concurrent increases in catabolic markers like RANTES and MIG remain counterintuitive. These specific elevations might account for the temporal increase in female lateral degradation scores at 12 months, as well as the observation that male Tac1−/− scores only trended to be lower than WT scores at the 18-month mark. Crucially, these systemic alterations cannot explain the profound structural protection observed in the αCGRP-/-mice, reinforcing the idea that systemic serum profiles may not accurately mirror the localized microenvironment of the knee joint.

Flow cytometry analyses of the synovial tissue were performed to provide insight into the localized immunological microenvironment. The data revealed pronounced sex-, age-, and genotype-dependent differences, elucidating the specific influence of SP and αCGRP on immune cells during age-related knee OA. In young male mice, all knockout models (Tac1−/−, αCGRP-/-, and DKO) exhibited a reduced overall macrophage count compared to WT controls, accompanied by a trending decrease in M2 macrophage polarization. Interestingly, this effect was mitigated in female mice, where single knockouts of SP or αCGRP were insufficient to alter macrophage counts. Only female DKO mice demonstrated a reduction in macrophages at a young age, suggesting a robust compensatory mechanism between the two neuropeptides in females. With advancing age, WT macrophage populations dropped to match KO levels across the panel, which aligns with recent reports demonstrating a general age-related decline in murine joint macrophages [32]. Mechanistically, existing in vitro evidence demonstrates that macrophages express functional αCGRP receptors and that αCGRP promotes tissue-repairing M2-like polarization, whereas αCGRP deficiency actively promotes M1 and inhibits M2 polarization [33, 34]. The role of SP, however, appears highly context-dependent. While SP is known to trigger pro-inflammatory responses via NF-κB activation and can be produced by macrophages themselves, it is also capable of inducing M2 polarization through the PI3K/Akt/mTOR/S6kinase pathway [35–38]. Our in vivo data reflect this dynamic complexity, confirming that the absence of these neuropeptides disrupts early local macrophage homeostasis.

Significant sexual dimorphism and neuropeptide interactions were equally evident within the T cell compartment. Young female WT mice presented with notably high CD8+ T cell numbers compared to KO models, once again pointing toward an interaction between SP and αCGRP. By 18 months, female WT levels decreased to align with the KO baseline. In stark contrast, males exhibited a significant age-related increase in CD8+ T cells exclusively in the single αCGRP-/-strain. Because this increase was absent in male DKO mice, a functional interplay with SP is highly probable. While T cells possess functional αCGRP receptors, we did not observe the previously reported αCGRP-driven shift toward a Th2 phenotype in our KO models [39, 40]. Instead, when considered alongside the loss of anti-inflammatory M2 macrophage polarization, it is plausible that αCGRP deficiency generates a localized pro-inflammatory milieu that allows cytotoxic CD8+ T cells to survive longer. Crucially, our data indicate this specific mechanism is exclusive to males. Furthermore, the established roles of SP in T cell regulation are inherently contradictory, spanning both the promotion of T cell proliferation and lymphocyte attraction as well as the reduction of CD8+ T cell migratory and proliferative activity [36, 41]. This context-dependent duality clearly manifests in our joint tissue findings, further underscored by the observation that female SP KO mice exhibited an age-dependent increase in CD4+ T cells, whereas males showed no such differences.

When interpreting these findings, methodological limitations must be considered. First, the study relies on life-long, systemic genetic knockout models. These long-term deletions may trigger developmental or compensatory mechanisms that obscure direct, localized effects. Second, while systemic serum markers were comprehensively profiled, analyzing local synovial fluid concentrations would have provided a much more accurate representation of the active biological processes within the joint. Unfortunately, murine anatomy and extremely low synovial fluid volumes render consistent synovial fluid collection and analysis highly impractical. Finally, it must be acknowledged that murine models of age-related spontaneous OA-like processes possess fundamental biological and mechanical differences from human pathology, most notably their quadrupedal biomechanics, shorter lifespan, and the strict absence of menopause in female subjects.

In conclusion, this study establishes that the sensory neuropeptides SP and αCGRP act as critical modulators of spontaneous, age-related OA. The long-term genetic deletion of SP and αCGRP confers significant protection against age-related cartilage degradation. While the structural preservation observed in SP-deficient models can be mechanistically linked to systemic cytokine alterations – most notably a reduction in IL-23 and an elevation of IP-10 –these systemic serum profiles do not account for the robust protection seen in αCGRP knockout models. This underscores a stark divergence between systemic circulating markers and the localized microenvironment of the joint. Furthermore, the deficiency of these neuropeptides disrupts early local macrophage homeostasis and significantly alters T cell populations within the synovial tissue in a manner heavily dictated by age, genotype, and sex. Ultimately, these results emphasize that future diagnostic and therapeutic strategies for age-related joint degeneration must account for complex sexual dimorphism and prioritize the localized synovial microenvironment rather than relying solely on systemic profiles.

## Supporting information

Pann et al. Supplementary Data

## Acknowledgements

We thank Anja Pasoldt, Maren Hofmann, Arisha-Johanna Patt, Ariane Santos-Goncalves for their excellent technical assistance in animal experiments, histology, and histomorphometry.

## Declarations

### Ethics approval

#### Title of the animal project

Untersuchung zum Einfluss des sensiblen Nervensystems auf Veränderungen des osteoarthrotischen subchondralen Knochengewebes unter definierter mechanischer Belastung (Laufbandtraining)

The local authorities in Würzburg (Regierung von Unterfranken) approved all animal experiments.

#### Approval number

AZ 55.2-2532-2-1253

#### Date of approval

November 5, 2020

### Author Approvals

All authors have seen and approved the manuscript and it hasn’t been accepted or published elsewhere.

### Consent for publication

Not applicable

### Availability of data and materials

The datasets used and/or analyzed during the current study are available from the corresponding author on reasonable request.

### Contribution

Patrick Pann: Project administration, investigation, methodology, validation, formal analysis, data curation, visualization, writing (original draft, review, editing). Ravikumar Mayakrishnan: Investigation, methodology, validation, formal analysis of FACS measurements, writing (review, editing). Babak Moradi: Methodology (FACS), resources, writing (review, editing), funding acquisition. Brian Johnstone: writing (review, editing). Susanne Grässel: Conceptualization, project administration, resources, writing (review, editing), funding acquisition.

## Supplementary Figures

**Supplementary Figure 1: Flow cytometric gating hierarchy for myeloid and lymphoid lineages.** Following the initial discrimination of single and viable events, lineage-specific marker panels were applied. (A) The lymphoid compartment was isolated based on CD3 positivity, enabling the subsequent stratification of CD4+ and CD8+ T-cell subsets. CD4+ subpopulations were further discriminated into Th1 (IFN-γ+), Th2 (IL-4+), Th17 (IL17a+), and T_reg_ cells (CD25+ FoxP3+). (B) The myeloid compartment was defined by CD11b expression, from which subpopulations were further delineated into neutrophils (CD11b+ Ly6G+) and macrophages (CD11b+ F4/80+ Ly6C-low). Macrophages were further stratified in M1 (CD11C+ CD206-) and M2 (CD11c-CD206+) macrophages.

**Supplementary Figure 2. Systemic profiling of low-abundance osteoarthritis-associated factors across aging neuropeptide-deficient mice.** Serum marker concentrations of female (left) and male (right) WT and KO mice at 6, 12, and 18 months of age. Statistical analysis is using Kruskal-Wallis with Dunn’s test for multiple comparisons. * p < 0.05, ** p < 0.01, *** p < 0.001. N = 2-8.

**Supplementary Figure 3. Flow cytometric subpopulation analysis of synovial T helper cells and polarized macrophages.** Flow cytometry analysis shows the percentages of synovial leukocyte subpopulations in female (A, C, E, G, I, K) and male (B, D, F, H, J, L) WT and KO mice at 6 and 18 months of age. Statistical analysis is using two-way ANOVA with Tukey’s test for multiple comparisons. * p < 0.05, ** p < 0.01. N = 3-8.

## References

1. Grässel S, Muschter D. Peripheral Nerve Fibers and Their Neurotransmitters in Osteoarthritis Pathology. International journal of molecular sciences. 2017;18. 10.3390/IJMS18050931.

2. Li F-X-Z, Xu F, Lin X, Wu F, Zhong J-Y, Wang Y, et al. The Role of Substance P in the Regulation of Bone and Cartilage Metabolic Activity. Front Endocrinol (Lausanne). 2020;11:77. 10.3389/fendo.2020.00077.

3. Jenei-Lanzl Z, Grässel S, Schaible H-G, Straub RH. Sympathosensory interactions - Possible relevance in osteoarthritis. Osteoarthritis Cartilage. 2026;34:511–22. 10.1016/j.joca.2025.12.001.

4. Hildebrandt A, Dietrich T, Weber J, Günderoth MM, Zhou S, Fleckenstein FN, et al. The dual pro-inflammatory and bone-protective role of calcitonin gene-related peptide alpha in age-related osteoarthritis. Arthritis Res Ther. 2023;25:244. 10.1186/s13075-023-03215-3.

5. Pann P, Kalke P, Maier V, Schäfer N, Clausen-Schaumann H, Schilling AF, et al. Decoding the impact of exercise and αCGRP signaling on murine post-traumatic osteoarthritis progression. Arthritis Research & Therapy. 2025;27:129. 10.1186/s13075-025-03589-6.

6. Pann P, Kalke P, Maier V, Schäfer N, Clausen-Schaumann H, Schilling AF, et al. Exercise-Dependent effects of substance P deficiency on joint degeneration and inflammation in a surgical mouse model of osteoarthritis. Arthritis Res Ther. 2025;27:224. 10.1186/s13075-025-03693-7.

7. Rapp AE, Wolter A, Muschter D, Grässel S, Lang A. Impact of sensory neuropeptide deficiency on behavioral patterns and gait in a murine surgical osteoarthritis model. J Orthop Res. 2024. 10.1002/jor.25949.

8. Nawaz H, Umer M, Noordin S, Bertilson BC, Li J, Ahmed AS, et al. Synovial Neuronal Changes in Knee Joint Osteoarthritis. Open Journal of Rheumatology and Autoimmune Diseases. 2016;6:26–33. 10.4236/ojra.2016.62005.

9. Muschter D, Fleischhauer L, Taheri S, Schilling AF, Clausen-Schaumann H, Grässel S. Sensory neuropeptides are required for bone and cartilage homeostasis in a murine destabilization-induced osteoarthritis model. Bone. 2020;133:115181. 10.1016/j.bone.2019.115181.

10. Zimmer A, Zimmer AM, Baffi J, Usdin T, Reynolds K, König M, et al. Hypoalgesia in mice with a targeted deletion of the tachykinin 1 gene. Proc Natl Acad Sci U S A. 1998;95:2630–5. 10.1073/pnas.95.5.2630.

11. Lu JT, Son Y-J, Lee J, Jetton TL, Shiota M, Moscoso L, et al. Mice Lacking α-Calcitonin Gene-Related Peptide Exhibit Normal Cardiovascular Regulation and Neuromuscular Development. Molecular and Cellular Neuroscience. 1999;14:99–120. 10.1006/mcne.1999.0767.

12. Glasson SS, Chambers MG, Van Den Berg WB, Little CB. The OARSI histopathology initiative – recommendations for histological assessments of osteoarthritis in the mouse. Osteoarthritis and Cartilage. 2010;18:S17–23. 10.1016/j.joca.2010.05.025.

13. Jackson MT, Moradi B, Zaki S, Smith MM, McCracken S, Smith SM, et al. Depletion of Protease-Activated Receptor 2 but Not Protease-Activated Receptor 1 May Confer Protection Against Osteoarthritis in Mice Through Extracartilaginous Mechanisms. Arthritis & Rheumatology. 2014;66:3337–48. 10.1002/art.38876.

14. Motulsky HJ, Brown RE. Detecting outliers when fitting data with nonlinear regression – a new method based on robust nonlinear regression and the false discovery rate. BMC Bioinformatics. 2006;7:123. 10.1186/1471-2105-7-123.

15. Schäfer N, Kalke P, Mayakrishnan R, Mei J, Kollatz L, Ehrnsperger M, et al. The α-MSH-MC1R Axis Modulates Sex-Specific Senescence and Inflammation Processes in Human Articular Chondrocytes and Mice Knee Joints. Aging and disease. 2026;:0. 10.14336/AD.2025.1307.

16. Li C, Zheng Z. Cartilage Targets of Knee Osteoarthritis Shared by Both Genders. Int J Mol Sci. 2021;22:569. 10.3390/ijms22020569.

17. Patel J, Chen S, Katzmeyer T, Pei YA, Pei M. Sex-dependent variation in cartilage adaptation: from degeneration to regeneration. Biol Sex Differ. 2023;14:17. 10.1186/s13293-023-00500-3.

18. Geraghty T, Obeidat AM, Ishihara S, Wood MJ, Li J, Lopes EBP, et al. Age-Associated Changes in Knee Osteoarthritis, Pain-Related Behaviors, and Dorsal Root Ganglia Immunophenotyping of Male and Female Mice. Arthritis Rheumatol. 2023;75:1770–80. 10.1002/art.42530.

19. Poudel SB, Ruff RR, Yildirim G, Miller RA, Harrison DE, Strong R, et al. Development of primary osteoarthritis during aging in genetically diverse UM-HET3 mice. Arthritis Res Ther. 2024;26:118. 10.1186/s13075-024-03349-y.

20. Jiang S, Xie W, Knapstein PR, Donat A, Albertsen L-C, Sevecke J, et al. Transcript-dependent effects of the CALCA gene on the progression of post-traumatic osteoarthritis in mice. Commun Biol. 2024;7:223. 10.1038/s42003-024-05889-0.

21. Lee KM-C, Lupancu T, Achuthan AA, De Steiger R, Hamilton JA. IL-23p19 in osteoarthritic pain and disease. Osteoarthritis and Cartilage. 2024;32:1413–8. 10.1016/j.joca.2024.05.011.

22. Askari A, Naghizadeh MM, Homayounfar R, Shahi A, Afsarian MH, Paknahad A, et al. Increased Serum Levels of IL-17A and IL-23 Are Associated with Decreased Vitamin D3 and Increased Pain in Osteoarthritis. PLoS ONE. 2016;11:e0164757. 10.1371/journal.pone.0164757.

23. Lee KM-C, Zhang Z, Achuthan A, Fleetwood AJ, Smith JE, Hamilton JA, et al. IL-23 in arthritic and inflammatory pain development in mice. Arthritis Res Ther. 2020;22:123. 10.1186/s13075-020-02212-0.

24. Najm A, McInnes IB. IL-23 orchestrating immune cell activation in arthritis. Rheumatology (Oxford). 2021;60 Suppl 4:iv4–15. 10.1093/rheumatology/keab266.

25. Donat A, Xie W, Jiang S, Brylka LJ, Schinke T, Rolvien T, et al. Cxcl9-deficiency attenuates the progression of post-traumatic osteoarthritis in mice. Inflamm Res. 2025;74:48. 10.1007/s00011-025-02013-8.

26. Bonfante HDL, Almeida CDS, Abramo C, Grunewald STF, Levy RA, Teixeira HC. CCL 2, CXCL 8, CXCL 9 and CXCL 10 serum levels increase with age but are not altered by treatment with hydroxychloroquine in patients with osteoarthritis of the knees. Int J of Rheum Dis. 2017;20:1958–64. 10.1111/1756-185X.12589.

27. Nees TA, Rosshirt N, Zhang JA, Reiner T, Sorbi R, Tripel E, et al. Synovial Cytokines Significantly Correlate with Osteoarthritis-Related Knee Pain and Disability: Inflammatory Mediators of Potential Clinical Relevance. J Clin Med. 2019;8:1343. 10.3390/jcm8091343.

28. Furman BD, Kent CL, Huebner JL, Kraus VB, McNulty AL, Guilak F, et al. CXCL10 is upregulated in synovium and cartilage following articular fracture. J Orthop Res. 2018;36:1220–7. 10.1002/jor.23735.

29. Agere SA, Akhtar N, Watson JM, Ahmed S. RANTES/CCL5 Induces Collagen Degradation by Activating MMP-1 and MMP-13 Expression in Human Rheumatoid Arthritis Synovial Fibroblasts. Front Immunol. 2017;8:1341. 10.3389/fimmu.2017.01341.

30. Takebe K, Rai MF, Schmidt EJ, Sandell LJ. The chemokine receptor CCR5 plays a role in post-traumatic cartilage loss in mice, but does not affect synovium and bone. Osteoarthritis and Cartilage. 2015;23:454–61. 10.1016/j.joca.2014.12.002.

31. Li F, Venkatesan JK, Madry H, Cucchiarini M. Therapeutic strategies targeting synovial cells to treat osteoarthritis. Biomedicine & Pharmacotherapy. 2025;189:118317. 10.1016/j.biopha.2025.118317.

32. Dapas M, DeJong EN, Wang Y, Mills C, Dowling SD, Mayer ML, et al. Sex-associated transcriptional changes to synovial macrophages in the aging joint. Front Immunol. 2026;17:1724385. 10.3389/fimmu.2026.1724385.

33. Asahina A, Moro O, Hosoi J, Lerner EA, Xu S, Takashima A, et al. Specific induction of cAMP in Langerhans cells by calcitonin gene-related peptide: relevance to functional effects. Proc Natl Acad Sci USA. 1995;92:8323–7. 10.1073/pnas.92.18.8323.

34. Yuan Y, Jiang Y, Wang B, Guo Y, Gong P, Xiang L. Deficiency of Calcitonin Gene-Related Peptide Affects Macrophage Polarization in Osseointegration. Front Physiol. 2020;11:733. 10.3389/fphys.2020.00733.

35. Sun J, Ramnath RD, Zhi L, Tamizhselvi R, Bhatia M. Substance P enhances NF-κB transactivation and chemokine response in murine macrophages via ERK1/2 and p38 MAPK signaling pathways. American Journal of Physiology-Cell Physiology. 2008;294:C1586–96. 10.1152/ajpcell.00129.2008.

36. O’Connor TM, O’Connell J, O’Brien DI, Goode T, Bredin CP, Shanahan F. The role of substance P in inflammatory disease. Journal Cellular Physiology. 2004;201:167–80. 10.1002/jcp.20061.

37. Hua F, Wang H-R, Bai Y-F, Sun J-P, Wang W-S, Xu Y, et al. Substance P promotes epidural fibrosis via induction of type 2 macrophages. Neural Regeneration Research. 2023;18:2252–9. 10.4103/1673-5374.369120.

38. Lim JE, Chung E, Son Y. A neuropeptide, Substance-P, directly induces tissue-repairing M2 like macrophages by activating the PI3K/Akt/mTOR pathway even in the presence of IFNγ. Sci Rep. 2017;7:9417. 10.1038/s41598-017-09639-7.

39. Levite M. Neurotransmitters activate T-cells and elicit crucial functions via neurotransmitter receptors. Current Opinion in Pharmacology. 2008;8:460–71. 10.1016/j.coph.2008.05.001.

40. Mikami N, Matsushita H, Kato T, Kawasaki R, Sawazaki T, Kishimoto T, et al. Calcitonin Gene-Related Peptide Is an Important Regulator of Cutaneous Immunity: Effect on Dendritic Cell and T Cell Functions. The Journal of Immunology. 2011;186:6886–93. 10.4049/jimmunol.1100028.

41. Strell C, Sievers A, Bastian P, Lang K, Niggemann B, Zänker KS, et al. Divergent effects of norepinephrine, dopamine and substance P on the activation, differentiation and effector functions of human cytotoxic T lymphocytes. BMC Immunol. 2009;10:62. 10.1186/1471-2172-10-62.

