## Supplementary material for "Sensory Neuropeptides Dictate Sex-Specific Synovial Immunity and Cartilage Degeneration in Aging Mice": Pann et al. Supplementary Data

**Affiliations**

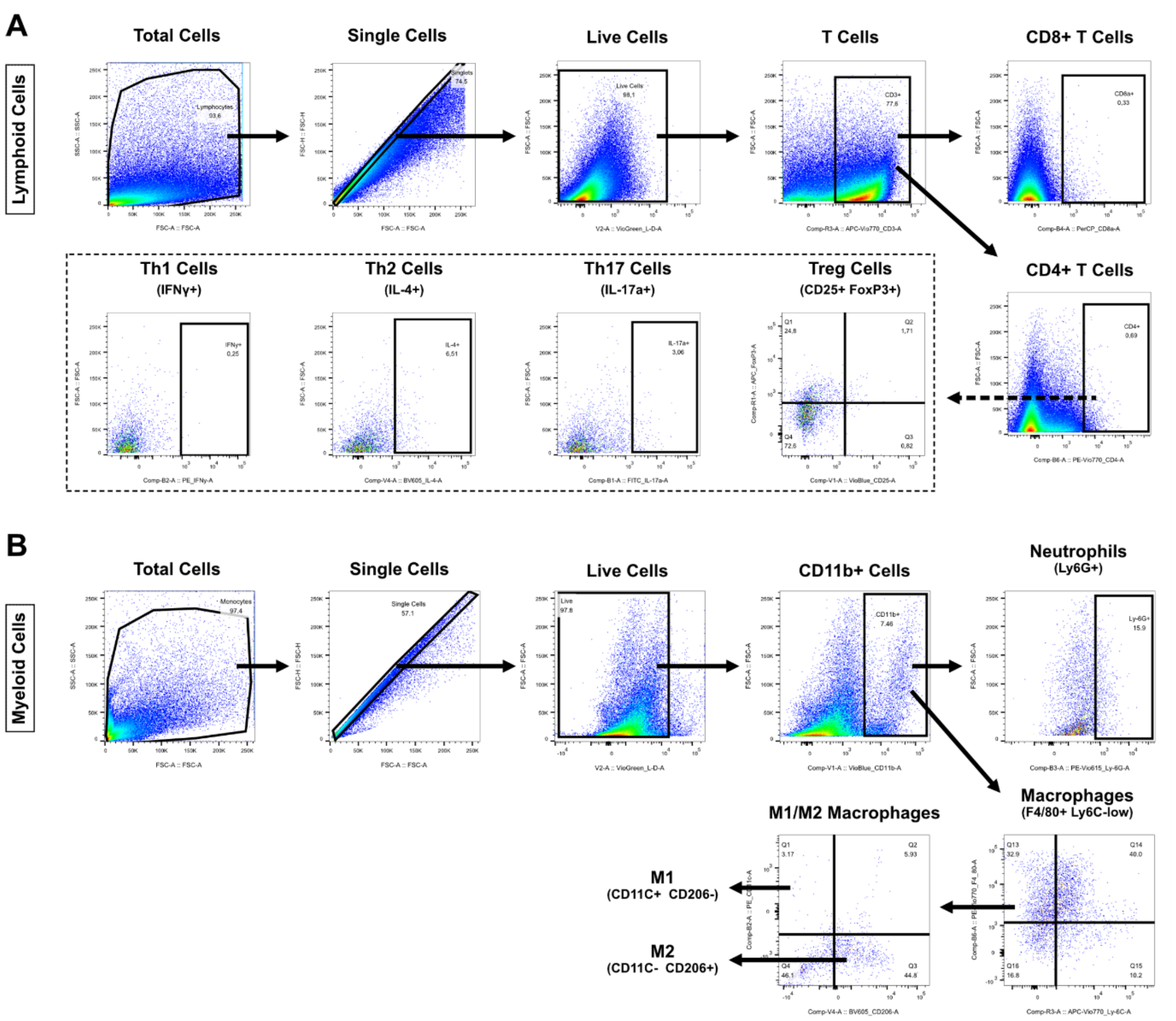


**Supplementary Figure 1: Flow cytometric gating hierarchy for myeloid and lymphoid lineages.**

Following the initial discrimination of single and viable events, lineage-specific marker panels were applied. (A) The lymphoid compartment was isolated based on CD3 positivity, enabling the subsequent stratification of CD4+ and CD8+ T-cell subsets. CD4+ subpopulations were further discriminated into Th1 (IFN-γ+), Th2 (IL-4+), Th17 (IL17a+), and $T_{reg}$ cells (CD25+ FoxP3+). (B) The myeloid compartment was defined by CD11b expression, from which subpopulations were further delineated into neutrophils (CD11b+ Ly6G+) and macrophages (CD11b+ F4/80+ Ly6C-low). Macrophages were further stratified in M1 (CD11C+ CD206-) and M2 (CD11c- CD206+) macrophages.

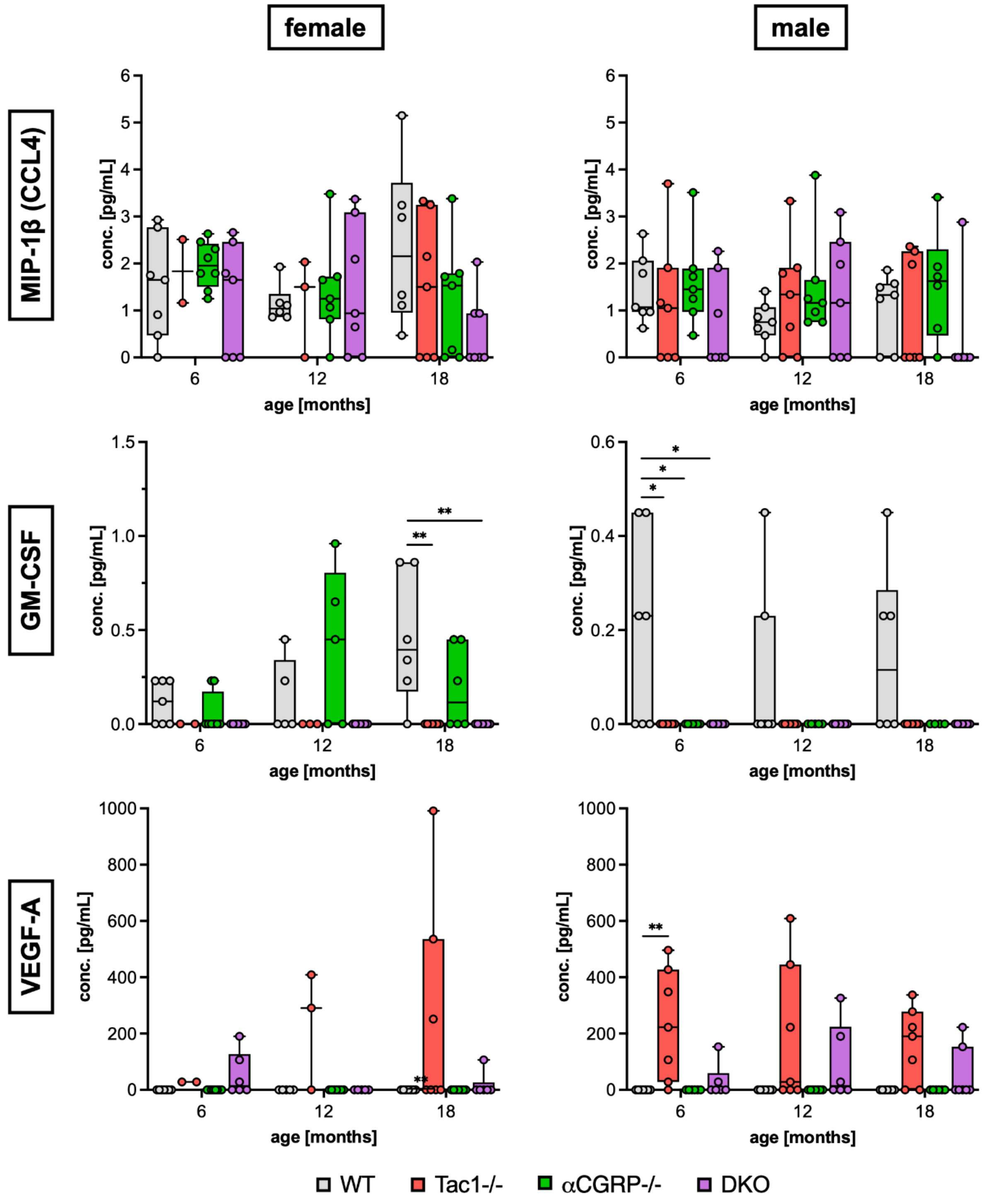


**Supplementary Figure 2. Systemic profiling of low-abundance osteoarthritis-associated factors across aging neuropeptide-deficient mice.**

Serum marker concentrations of female (left) and male (right) WT and KO mice at 6, 12, and 18 months of age. Statistical analysis is using Kruskal-Wallis with Dunn's test for multiple comparisons. * $p < 0.05$, ** $p < 0.01$, *** $p < 0.001$. N = 2-8.

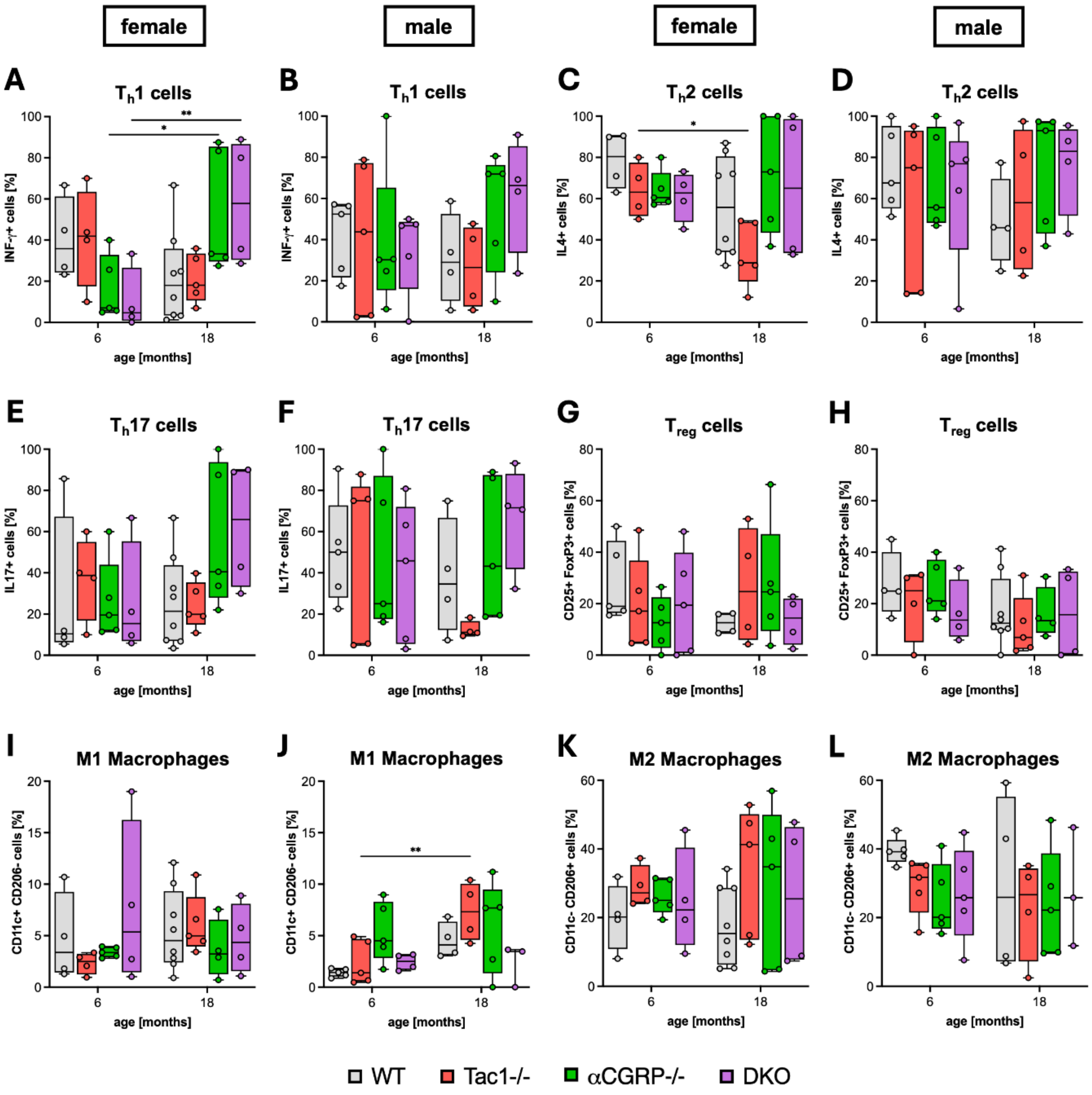


**Supplementary Figure 3. Flow cytometric subpopulation analysis of synovial T helper cells and polarized macrophages.**

Flow cytometry analysis shows the percentages of synovial leukocyte subpopulations in female (A, C, E, G, I, K) and male (B, D, F, H, J, L) WT and KO mice at 6 and 18 months of age. Statistical analysis is using two-way ANOVA with Tukey’s test for multiple comparisons. * $p < 0.05$, ** $p < 0.01$. N = 3-8.
